# Continent-scale phenotype mapping using citizen scientists’ photographs

**DOI:** 10.1101/503847

**Authors:** Jonathan P. Drury, Morgan Barnes, Ann E. Finneran, Maddie Harris, Gregory F. Grether

## Abstract

Field investigations of phenotypic variation in free-living organisms are often limited in scope owing to time and funding constraints. By collaborating with online communities of amateur naturalists, investigators can greatly increase the amount and diversity of phenotypic data in their analyses while simultaneously engaging with a public audience. Here, we present a method for quantifying phenotypes of individual organisms in citizen scientists’ photographs. We then show that our protocol for measuring wing phenotypes from photographs yields accurate measurements in two species of Calopterygid damselflies. Next, we show that, while most observations of our target species were made by members of the large and established community of amateur naturalists at iNaturalist.org, our efforts to increase recruitment through various outreach initiatives were successful. Finally, we present results from two case studies: (1) an analysis of wing pigmentation in male smoky rubyspots (*Hetaerina titia*) showing previously undocumented geographical variation in a seasonal polyphenism, and (2) an analysis of variation in the relative size of the wing spots of male banded demoiselles (*Calopteryx splendens*) in Great Britain questioning previously documented evidence for character displacement. Our results demonstrate that our protocol can be used to create high quality phenotypic datasets using citizen scientists’ photographs, and, when combined with metadata (e.g., date and location), can greatly broaden the scope of studies of geographical and temporal variation in phenotypes. Our analyses of the recruitment and engagement process also demonstrate that collaborating with an online community of amateur naturalists can be a powerful way to conduct hypothesis-driven research aiming to elucidate the processes that impact trait evolution at landscape scales.

## Introduction

Traditionally, studies of phenotypic geographic variation have been based on specimens collected by scientists, but due to time and funding constraints, the geographic coverage of such datasets is often rather limited. Studying geographic variation in traits that change over time (e.g., seasonal polyphenisms) can be especially challenging. The problems with relying on datasets of limited, and potentially biased, geographic and temporal coverage are well documented (Siepielski et al. 2009, Powers et al. 2011, Kramer-Schadt et al. 2013). Such limits can impact the reliability of inferences about the presence of lineages in a particular area (Isaac et al. 2014). For example, the proportion of records of threatened bird species in a dataset strongly depends on the source and time period of data collection (Boakes et al. 2010). Sampling biases may also impact inferences about geographic variation in phenotypes. For instance, though sympatric ‘shifts’ in bill length of rock nuthatches (*Sitta tephronota* & *S. neumayer*) were originally presented as evidence for character displacement between the two species (Vaurie 1951, Brown and Wilson 1956), subsequent analyses show that these shifts can be alternatively explained by a longitudinal cline in bill morphology (Grant 1972).

In the past decade, many investigators have increasingly engaged with lay people, often referred to as ‘citizen scientists’, to generate continental-scale datasets (Silvertown 2009, Dickinson et al. 2010, Sullivan et al. 2014, Irwin 2018). On many web platforms for amateur naturalists, users submit photographs of organisms they observe, along with the time and location of their observations. By facilitating coverage of a greater geographic area with greater temporal resolution, quantifying phenotypes from photographs taken by citizen scientists could offer a powerful alternative to traditional approaches.

One way that investigators can launch a citizen science initiative relatively easily is through iNaturalist (http://www.inaturalist.org), a web platform run by the California Academy of Sciences. With over 750k users, the iNaturalist community provides a powerful means to quantify broad-scale geographical and temporal patterns. Moreover, photographic observations are vetted by other users on the platform, meaning that observations pass quickly through a built-in quality-filtering step. Observations from the iNaturalist.org platform have made it possible for investigators to conduct wide-ranging analyses, such as monitoring range expansions (Mori et al. 2018), mapping public health risks for planning interventions (Geneviève et al. 2018), and identifying hotspots of bird mortality due to window collisions (Winton et al. 2018). Here, we describe a method for launching a citizen scientist initiative through iNaturalist.org and generating accurate phenotypic measurements from users’ photographs. Using this approach, we then present the results from two case studies, in which we test hypotheses about temporal and geographic variation in the wing colour of males from two species of Calopterygid damselflies (*Hetaerina titia* and *Calopteryx splendens*).

## Materials and Methods

### Generating phenotypic datasets from amateur naturalists’ photographs

To assess the effectiveness of generating phenotypic datasets from amateur naturalists’ photographs, we launched citizen science initiatives to collect photographs of three species of Calopterygid damselflies: smoky rubyspots (*Hetaerina titia*; Fig. 1c,e), banded demoiselles (*Calopteryx splendens*; Fig. 1d,f), and beautiful demoiselles (*Calopteryx virgo*). We used iNaturalist to solicit observations from amateur naturalists, tracking the impact of our recruitment strategies on observation accrual throughout. In addition, we conducted fieldwork to develop and validate a measurement pipeline for extracting data from the photographs submitted to iNaturalist by amateur naturalists.

**Figure 1.**
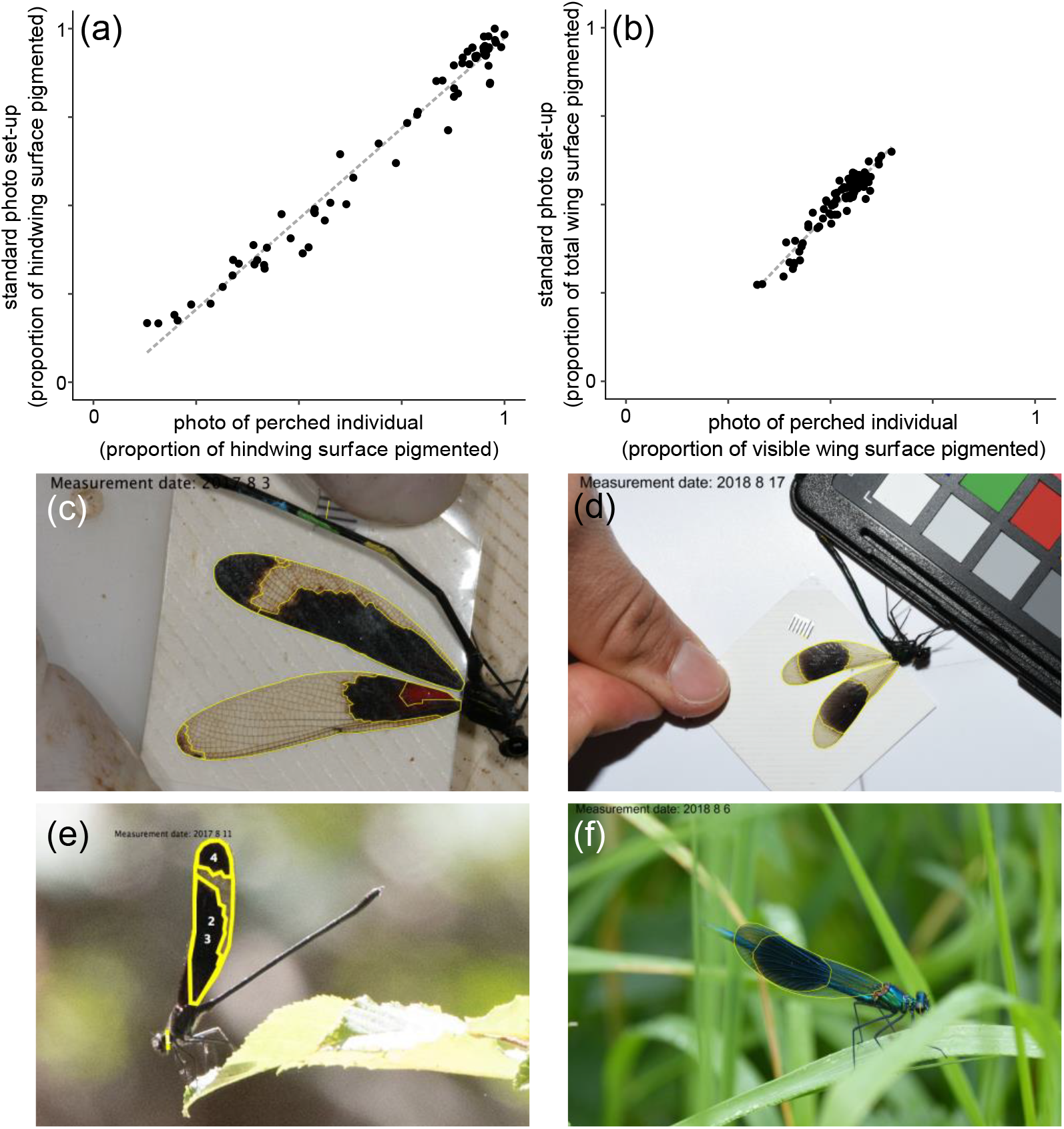
Measurements of relative wing pigmentation extracted from photos of perched individuals are highly correlated with measurements extracted from photos taken under standardised conditions for both (a) smoky rubyspots (*H. titia*) (presenting hindwing measurements, which were used in analyses in Case Study #1) and (b) banded demoiselles (*C. splendens*) (presenting total wing measurements, which were used in analyses in Case Study #2). Examples of photos taken under standardised and perched conditions are shown for (c,e) *H. titia* and (d,f) *C. splendens*. Yellow lines represent measurements made on each photo.

The website iNaturalist.org allows users to create a “project page”, where either citizen scientists can directly submit photographs or project managers can attach existing iNaturalist observations. We created two such project pages: one, created in July 2017, for smoky rubyspot damselflies throughout their range in North America (https://www.inaturalist.org/projects/smoky-rubyspots), and another for banded and beautiful demoiselles in the United Kingdom, created in May 2018 (https://www.inaturalist.org/projects/british-demoiselles). For both projects, we regularly searched iNaturalist for observations that we could add to each project page until 5 November 2018. In total, we compiled 620 observations of smoky rubyspot damselflies, 409 observations of banded demoiselles, and 120 observations of beautiful demoiselles during these time periods.

Male smoky rubyspots and banded demoiselles possess discrete patches of melanised pigment on their wings that influence their territorial and mating interactions (Tynkkynen et al. 2006, Svensson et al. 2007, Anderson and Grether 2010, Drury and Grether 2014), so we focused on quantifying variation in the size of these patches relative to the entire wing surface. Beautiful demoiselles have nearly entirely melanised wings, so while we included data from beautiful damselflies in our analyses of observation rates on iNaturalist, we limited our analyses of wing phenotypes to male smoky rubyspots and male banded demoiselles.

To determine if measurements collected from citizen scientists’ photos are reliable and accurate, we also conducted a field study to compare measurements of the proportion of wing surfaces with melanised pigment taken under standardised settings to measurements extracted from photographs of perched males (i.e., photographs similar to those submitted to iNaturalist). For both species included in our analyses (*H. titia* & *C. splendens*), we visited several streams (Supplementary Table A1) and walked along the bank in search of perched individuals. When we encountered perched individuals, we photographed them (using a Nikon D750 for *H. titia* and Panasonic Lumix DMC-FZ60 for *C. splendens*), then captured them and marked their abdomen with a unique colour combination (Anderson et al. 2011) so that we could later match the perched photograph with a photograph taken under standard conditions.

To test whether we could obtain reliable, accurate wing phenotype data from photos taken of perched individuals, we quantified the proportion of melanised pigment on the wings of mature males using photos taken under standard conditions. That is, we photographed the left forewing and hindwing of individuals (using Canon EOS 20D with a 100 mm macro lens for *H. titia* and a Nikon D7200 with an 18-55mm zoom lens for *C. splendens*) on a level stage with a millimetre scale in the same plane as the wings.

For both perched and standard photographs, we measured each wing manually using NIH ImageJ and a custom macro script (Supplementary material Appendix 1, 2). For photographs of perched individuals, we took two approaches: (1) outlining the entire visible wing surface and all the black pigment that was visible to generate a measurement of the ‘entire wing proportion melanised’, or the total visible area of black pigment divided by the total visible area of the wing, and (2) carefully isolating and outlining the hindwing surface and the hindwing pigmented regions, to measure the ‘hindwing proportion melanised’ (Supplementary material Appendix 1). For the standard photographs, we measured wing surface area and pigmented area separately on the left forewing and hindwing, which generated two measurements: (1) the ‘total proportion melanised’, or the relative area of both the forewing and hindwing surface covered in black pigment, and (2) the ‘hindwing proportion melanised’ (which might be more comparable to measurements from photos of perched damselflies, as the hindwings cover forewings when damselflies are perched). We then compared the measurements from standard and perched photographs using linear regressions.

For both species, we measured the wings and wing spots of citizen science photographs as described for perched individuals above. In some cases, we were able to measure the hindwing in a photo from a citizen scientist, but not the entire wing surface, or vice-versa. In these instances, given that the two measurement types were highly correlated (Adjusted R^2^ for *C. splendens* = 0.93; *H. titia* = 0.99), we used predictions from linear regression to interpolate missing data where possible.

### Recruitment strategies

We took different approaches to recruiting observations for the two projects. For the smoky rubyspot project, we added observations to the project page from July 2017-November 2018. Although our web page contains instructions for taking high quality photographs and is publicly visible to iNaturalist users, we did not otherwise take any measures to actively recruit citizen scientists to the project after its initial launch. For the demoiselle project, created in May 2018, we actively recruited participants in two ways. First, we contacted the British Dragonfly Society, which advertised our project and invited participants to submit photographs in a monthly newsletter sent out to members. Second, we systematically monitored posts to http://www.twitter.com for photographs of demoiselles in the U.K., and asked Twitter users to upload their photographs on iNaturalist.

To determine whether our recruitment efforts influenced the rate of citizen scientist participation, we conducted three sets of analyses of the accrual of observations of males and females of each of the three species in our projects. First, for each project separately, we compared the number of observations per month before and after the project launch. Since the overall accrual rate of observations on iNaturalist has increased rapidly through time (Irwin 2018), we compared the count of observations of the focal species with that of a relevant “benchmark” group (sensu Hill 2012, see also Maes and Swaay 1997). For the smoky rubyspot project, this was other *Hetaerina* spp. within 50km of a site where *H. titia* was observed; for the British demoiselle project, this was all other damselflies observed within 50km of a site where *Calopteryx* spp. were observed. To account for variation across the year in the rate of upload (e.g., due to seasonality in species occurrence), we fit a mixed-effect Poisson regression model to the count data with a random intercept of the month of the observations using the R package lme4 (Bates et al. 2015) in R. For these models, since our main hypothesis test was whether the accrual of new observations for the focal taxon outpaced that of the benchmark taxa, we compared the fit of models with an interaction between the effect of the post-launch period and the focal taxon on observation accrual to models without that interaction.

For our third analysis, we directly compared whether the recruitment strategy for each species influenced the rate of observations. To do this, we first created a list of users who submitted observations of either the focal species or the relevant comparison group. Among these users, we considered individuals whose first observations in our dataset were of a focal species after the project launch date to be ‘newly-recruited’ users. This approach allowed us to monitor recruitment from within the iNaturalist platform (e.g., of users who had previously not submitted observations of damselflies, in the case of the British demoiselle project). We then tested whether the count of observations from newly recruited users differed between the two projects, fitting a mixed-effect Poisson regression model as described above. In this analysis, we compared the fit of models with an interaction between the project ID and whether observations came from new recruits to a model without the interaction. We also tested whether the relative number of users newly recruited to the iNaturalist platform differed across projects.

### Case study #1: Smoky rubyspot damselflies

Smoky rubyspots (*Hetaerina titia*) are distributed throughout North America, from Costa Rica in the south to Ontario in the north. Previous research has documented a seasonal polyphenism in several populations in drainages that empty into the Gulf of Mexico (Drury et al. 2015). In these Atlantic populations, males and females exhibit light-phase phenotypes that resemble other, sympatric congeners at the beginning of the breeding season; but, as the season progresses, the adults emerging have increasingly melanised wings, until dark-phase phenotypes comprise the majority. The striking difference between light-phase and dark-phase phenotypes led some investigators to conclude that they represent separate species (Johnson 1963), though they are currently classified as one (Garrison 1990, Drury et al. 2015). Our own analyses of populations on the Pacific slope of Central America (unpublished data) and preliminary observations of citizen scientists’ photographs from the Northern U.S. and Ontario suggest that the polyphenism is reduced or absent in some populations. Additionally, owing to our own time constraints and research goals, our previous research did not extend beyond the peak breeding season. Thus, the complete annual trajectory of the seasonal polyphenism, and the way in which it varies geographically, has remained unknown.For smoky rubyspots, we found that the relative wing pigmentation measurement from the isolated hindwing was the most reliable (see Results), so we focused our subsequent analyses on this measurement. However, results were qualitatively identical using the other measurement (see Supplementary material Table A4). To test the hypothesis, drawn from our personal observations in the field and preliminary observations of online records, that the seasonal polyphenism proceeds differently in Pacific and Northern populations from the way that was documented previously in Atlantic populations, we fit linear mixed-effect models in lme4 where the proportion of the hindwing with pigment varied according to the region an observation came from. These regions were identified a priori via visual inspection of observations on a map, with Northern populations being those which were separated by a large distance from populations contiguous with Gulf Coast populations (minimum distance = 290km), and Pacific populations being those in drainages that empty into the Pacific Ocean. We also included a quadratic function of observation date and latitude and longitude as fixed effects in the model. To account for potential pseudoreplication arising from multiple observations submitted at the same place, we created a ‘site’ term to aggregate observations from within 1.11 km (0.01 decimal degrees) (Kuussaari and Helio 2007, Roy et al. 2012). We then fit linear mixed effect models with a random effect of ‘site’ using the R package lme4 (Bates et al. 2015). We verified that spatial autocorrelation in our data did not affect our analyses by calculating Moran’s I on the model residuals using the R package ape (Paradis 2011). To test the hypothesis that patterns of wing pigmentation differ across the regions described above, we compared models with the ‘region’ variable to those without the ‘region’ variable. In addition, we fit a model where the Atlantic/Pacific split is preserved but with the northern and southern Atlantic regions pooled. Finally, we conducted an analysis on binned monthly observations at each site (i.e., where the response variable was the mean wing phenotype present each month), adding the number of observations in each bin as a predictor variable, to check whether our conclusions are impacted by variation in observation rates.

### Case study #2: Banded demoiselles

Banded and beautiful demoiselles have broadly overlapping ranges in the Western Palearctic. Beautiful demoiselles (*Calopteryx virgo*) have nearly solid black wings, while banded demoiselles (*C. splendens*) have transparent wings with a dark wing spot in the centre of each wing. Owing to their phenotypic and ecological similarity, combined with their broad geographic overlap, these species have become a model system for testing hypotheses about the impact of species interactions on trait evolution and coexistence. For instance, there is evidence throughout much of their range that character displacement has acted on banded demoiselle wing spot size, resulting in smaller wing spots where the two species are sympatric (Tynkkynen et al. 2004, Honkavaara et al. 2011) and consequently, lower levels of behavioural interference (Tynkkynen et al. 2005, 2006, Svensson et al. 2007, 2010). In Great Britain, there is evidence of a sympatric shift in male wing spot size (Honkavaara et al. 2011). However, there is also evidence for an effect of latitude (Hassall and Thompson 2009), longitude (Upton et al. 2016), and sampling date (Upton et al. 2016) on wing spot size in Great Britain. Until now, no study has attempted to disentangle the impact of latitude, longitude, date, and sympatry on the wing spots of banded demoiselles. Specifically, once the known impact of latitude, longitude, and date are accounted for, is there still evidence that character displacement has impacted wing pigmentation in male banded demoiselles living in sympatry with beautiful demoiselles?

For banded demoiselles, we conducted multiple regressions to identify the relative contribution of observation date, sympatry with *C. virgo*, latitude, and longitude on the relative size of the wing spot in males. We found that the relative wing pigmentation measurement from the entire wing surface was the most reliable measurement for *C. splendens* (see Results), and thus used these measurements in downstream analyses, though results were qualitatively identical using measurements isolated from the hindwing (see Supplementary Material Table A7). To identify observations from sympatric sites, we first downloaded observations of *C. virgo* from the past decade (2008 onward) from the Global Biodiversity Information Facility (GBIF.org 2018). We then calculated (1) the distance between the closest such observation and a given iNaturalist submission of *C. splendens* and (2) the distance from a *C. splendens* observation at which five records of *C. virgo* occur. We considered observations of *C. splendens* that were within 20km of the nearest *C. virgo* observation and 40km of the nearest 5 *C. virgo* observations to be sympatric. The resulting designations closely correspond to zones of range overlap indicated in field guides (Supplementary Material Fig. A1). As described above for *H. titia*, we created a ‘site’ term to aggregate proximate observations, fit linear mixed effect models, and verified that residuals were not spatially autocorrelated. To test for character displacement, we compared the fit of a model with a term indicating sympatry with *C. virgo* to a model without this sympatric term. To evaluate whether our conclusions were affected by the threshold we used to designate populations as sympatric, we repeated analyses using two additional thresholds with more conservative indices of whether banded demoiselle populations were sympatric with beautiful demoiselles. Additionally, we conducted an analysis on binned monthly observations at each site (as above), to evaluate whether variation in observation rates affected the results.

## Results

### Validation analyses

For both smoky rubyspots and banded demoiselles, the standardized measurements of the proportion of the wings that are pigmented are highly correlated with the measurements made from photographs of perched males (Fig. 1, Supplementary Material Table A2). Moreover, measurements within each class (i.e., standard or perched males) are highly reliable across observers (Supplementary material Table A3). The relative area of hindwing pigmentation was slightly more reliable for *H. titia*, while the relative area of pigment on the entire wing surface was slightly more reliable for *C. splendens* (Supplementary material Table A2), so we used the more reliable patches for each species in our subsequent analyses of citizen scientists’ photos.

### Effects of recruitment strategies

While the number of observations increased after the project launch date for both projects, the accrual of new observations outpaced that of the relevant benchmark group only for the demoiselle project (Fig. 2a,b; Table 1). For both projects, most observations were generated by the existing iNaturalist.org community, though recruitment efforts for the demoiselle project did successfully recruit new users to the iNaturalist platform: 10.4% [32/307] of users who ever submitted a damselfly observation in the U.K. joined iNaturalist after the project launch date and submitted a demoiselle as their first observation, compared to only 0.002% [2/1154] of users who ever submitted a rubyspot observation that joined iNaturalist after the project launch and submitted a smoky rubyspot as their first observation (Fisher’s exact test *p* < 0.0001).

**Figure 2.**
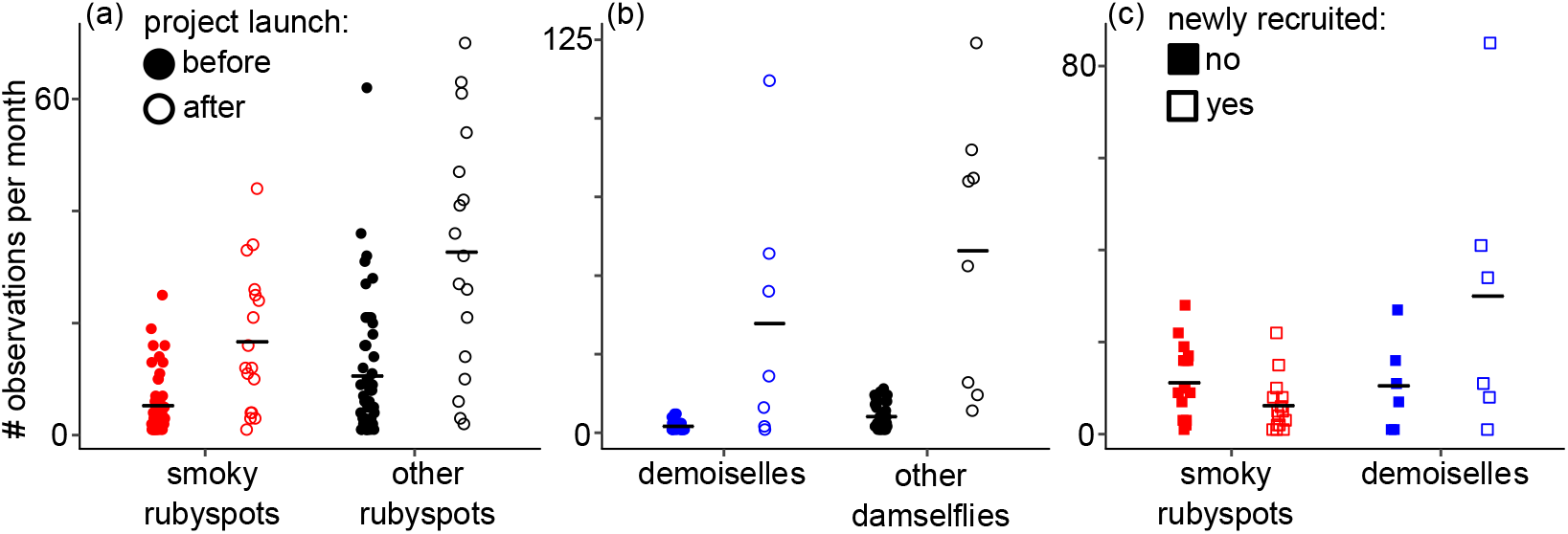
Effect of the active recruitment effort for the British demoiselle project, compared with the passive approach employed for the smoky rubyspot project [see Methods]. (a-c) While the overall rate of observations increased after the project launch, only with the demoiselles did this outpace the rate of observations of a relative comparison group (other rubyspot damselflies, in the case of the smoky rubyspot project, and all other damselflies, in the case of the British demoiselle project). (c) Newly-recruited users contributed a higher number of demoiselle observations after the project launch, compared with smoky rubyspot observations.

**Table 1.** Regression fits for analyses of mixed effect Poisson models of the monthly number of observations, with month included as a random effect. Analysis 1 compares the accrual of observations for smoky rubyspots (*n* = 546; 67 month-level observations) vs. benchmark taxa (*n* = 1125; 71 month-level observations), analysis 2 compares the accrual of observations for demoiselle (n = 300; 35 month-level observations) vs. benchmark taxa (*n* = 647; 44 month-level observations), and analysis 3 compares the post-launch accrual of observations for the smoky rubyspot (*n* = 283; 32 month-level observations) and demoiselle (*n* = 243; 12 month-level observations) projects. The 95% confidence intervals for terms presented in bold (calculated from the likelihood profile, using the ‘profile’ method in the confint function in lme4) do not cross 0. ΔAIC for model with interaction are calculated against a model without the interaction term (AIC_[model with interaction]_ – AIC_[model without interaction]_). “Focal taxon” refers to smoky rubyspots in analysis 1 and demoiselles in analysis 2. “After launch” refers to observations accrued after the project launch date.

| analysis | term | estimate | confidence intervals |  |
| --- | --- | --- | --- | --- |
|  |  |  | lower | upper |
| 1. smoky rubyspot project<br>vs. other rubyspots<br>$\Delta AIC = 1.97$ | **intercept** | **2.16** | **1.79** | **2.53** |
|  | **focal taxon** | **-0.66** | **-0.80** | **-0.51** |
|  | **after launch** | **1.12** | **1.00** | **1.24** |
|  | focal taxon * after launch | -0.02 | -0.22 | 0.19 |
| 2. British demoiselle project<br>vs. other damselflies<br>$\Delta AIC = -2.27$ | **intercept** | **1.20** | **0.68** | **1.71** |
|  | **focal taxon** | **-0.93** | **-1.24** | **-0.64** |
|  | **after launch** | **2.64** | **2.46** | **2.82** |
|  | **focal taxon \* after launch** | **0.35** | **0.02** | **0.69** |
| 3. smoky rubyspot project vs.<br>British demoiselle project<br>(post-launch observations)<br>$\Delta AIC = -81.92$ | **intercept** | **1.47** | **0.76** | **2.11** |
|  | **project = smoky rubyspot** | **0.47** | **0.19** | **0.77** |
|  | **new recruits** | **1.03** | **0.75** | **1.33** |
|  | **project\* new recruits** | **-1.70** | **-2.09** | **-1.33** |

Overall, the increased efforts to recruit iNaturalist users to the demoiselle project appear to have worked: there was an increased relative contribution of newly recruited users to the British demoiselle project, relative to the smoky rubyspot project (Fig. 2c; Table 1).

### Case study #1: Smoky rubyspots

Analyses of *H. titia* males show that the seasonal pattern of wing pigmentation differs across the three regions we analysed (Figures 3, A2, Tables 2, A4). In the northern regions, males tend to emerge much later in the year than in the other two regions, with later emerging individuals in this region having decreasing amounts of pigment on their wings. In the Atlantic region, where previous research documented the seasonal polyphenism (Drury et al. 2015), there is a large increase in the number of individuals with highly pigmented wing surfaces, and then a sharp decrease as the peak breeding season ends. We found a similarly cyclical pattern in the Pacific region, but the maximum level of pigmentation in this region was overall much lower than in the Atlantic region (Figure 3). We found that including the regions we generated based on our previous data in the model greatly improved the model fit (Tables 2, A5, Figure A2). We found no evidence that the number of observations affected our conclusions about spatial and temporal variation in wing pigmentation (Table A6).

**Figure 3.**
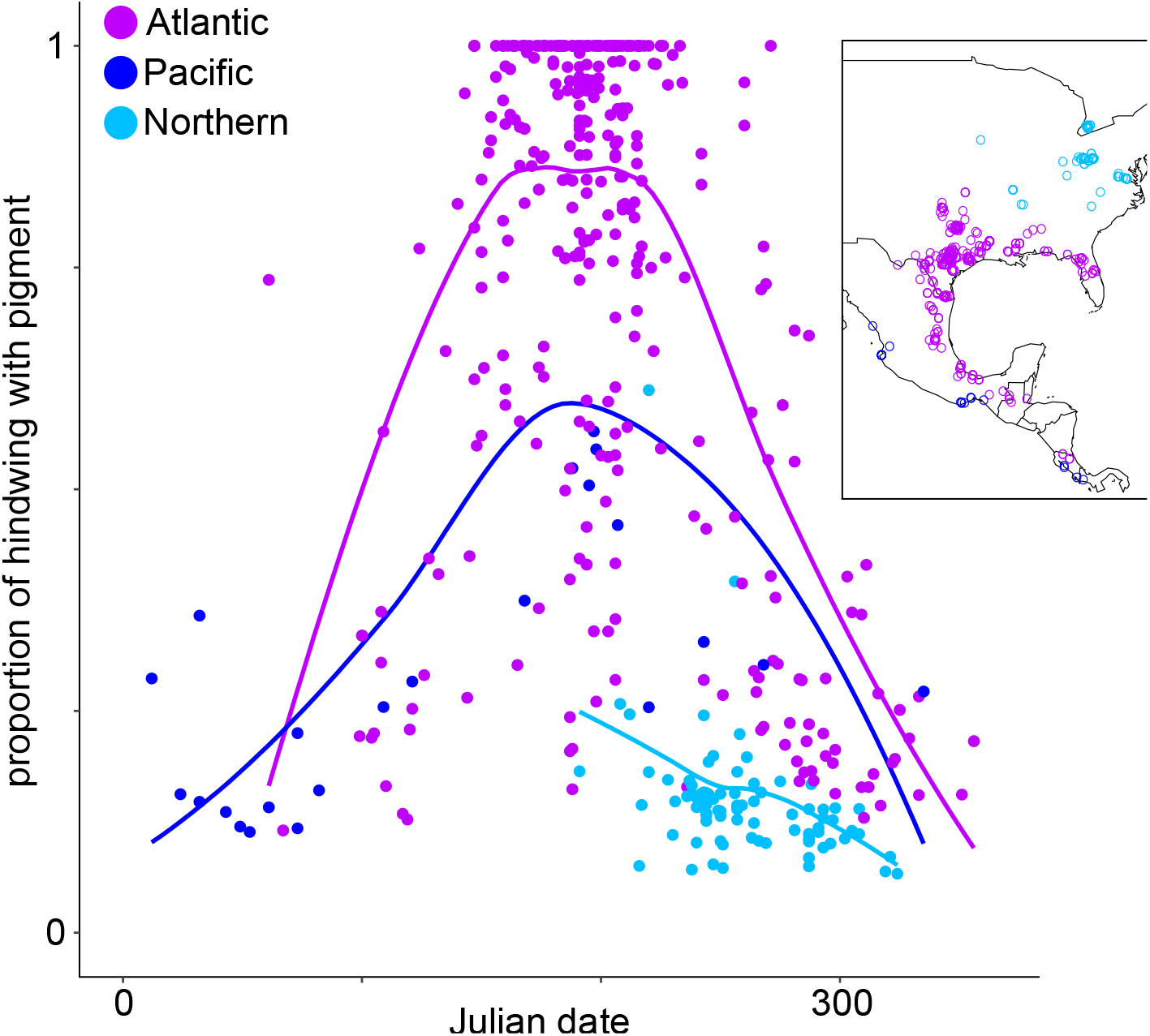
In Atlantic and Pacific populations of smoky rubyspots, there is a seasonal shift in the proportion of male wings with pigment, with dark-phase individuals predominating in the mid-season and light-phase forms predominating before and after the peak breeding season. Pacific populations exhibit a shallower polyphenism than Atlantic populations. Northern populations emerge later, and within these populations, later-emerging individuals tend to have proportionally less pigment on their wings. Inset: map of all iNaturalist observations of *H. titia*. Curves are plotted using a loess smoothing function on model predictions (using geom_smooth in the R-package ggplot2).

**Table 2.** Mixed-effect regression of the proportion of the hindwing surface with pigment in *H. titia* males. The dependent variable was log-transformed, and longitude, latitude, julian date and julian date were standardized to a mean of zero and variance of 1, prior to analysis. Samples with the same longitude and latitude (rounded to two decimal places) were grouped together under a random effects “site” term, to eliminate spatial autocorrelation (n = 437 samples, 271 groups), the absence of which was verified with Moran’s I (observed = −.0194, expected = −0.0026, sd = 0.0308, *p* = 0.58). The 95% confidence intervals (calculated from the likelihood profile, using the ‘profile’ method in the confint function in lme4) for terms presented in bold do not cross 0 (in this case, all variables in the model). A model excluding regional groupings provided a far worse fit to the data than the model presented here (AIC_[model with ‘region’]_ – AIC_[model without ‘region’]_ = −91.63; Table A5, Fig. A2). Tukey’s method was used for the multiple comparisons between regions.

**a. fixed effects**
| model term | estimate | confidence int. |  |
| --- | --- | --- | --- |
|  |  | lower | upper |
| intercept | **-0.60** | **-0.66** | **-0.53** |
| Northern region | **-1.10** | **-1.33** | **-0.86** |
| Pacific region | **-0.48** | **-0.76** | **-0.20** |
| Julian date | **1.55** | **1.33** | **1.76** |
| (Julian date)<sup>2</sup> | **-1.71** | **-1.92** | **-1.50** |
| latitude | **-0.08** | **-0.15** | **-0.01** |
| longitude | **-0.08** | **-0.15** | **-0.003** |

**b. multiple comparisons**
| contrast | estimate | std. error | z-value | p-value |
| --- | --- | --- | --- | --- |
| Northern-Atlantic | -1.10 | 0.12 | -9.03 | <0.001 |
| Pacific-Atlantic | -0.48 | 0.14 | -3.38 | 0.002 |
| Pacific-Northern | 0.62 | 0.21 | 2.88 | <0.001 |

### Case study #2: Banded demoiselles

After controlling for other geographic patterns, we did not find evidence for character displacement in the wing phenotypes of *C. splendens* males; males in sympatric populations do not have relatively smaller wing spots than males in allopatric populations (Tables 3, A7). We did, however, find an effect of observation date and latitude: males emerging earlier in the breeding season have relatively smaller wing spots than males emerging later in the season (Figure 4a), and males at higher latitudes tend to have relatively smaller wing spots (Figure 4b). These patterns were robust to different thresholds for designating sympatry (Table A8), and were not affected by variation in the number of observations across sites (Table A9).

**Figure 4.**
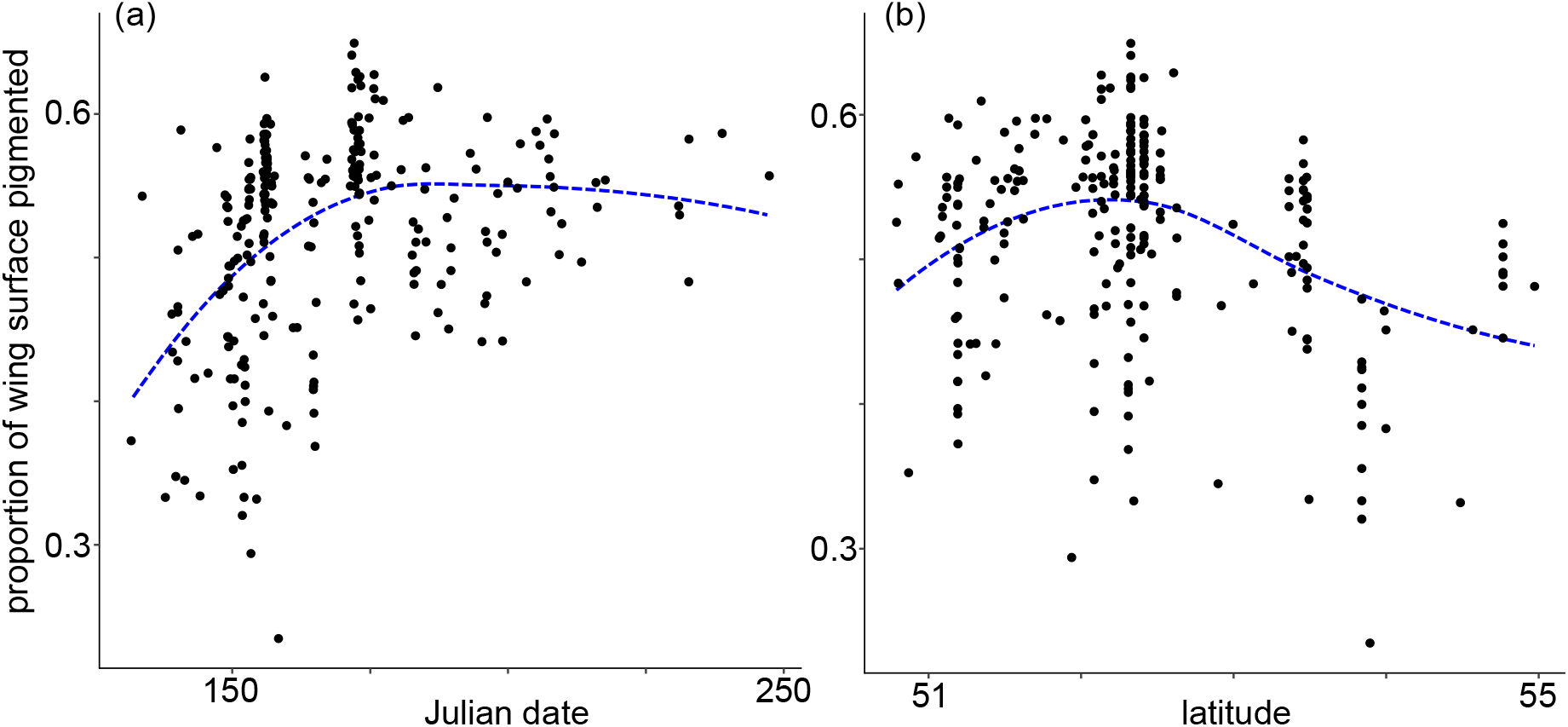
(a) The relative size of male banded demoiselle (*C. splendens*) wing spots in Great Britain increases with the date, such that early emerging individuals have relatively smaller wing spots. (b) Relative wing spot size also tends to decrease at higher latitudes. Curves are plotted using a loess smoothing function on model predictions (using geom_smooth in the R-package ggplot2).

**Table 3.** Mixed-effect regression of the log-transformed proportion of the visible wing surface with pigment for *C. splendens* males. Julian date and its quadratic term, latitude, and longitude were all z-transformed prior to model fitting. The 95% confidence intervals (calculated from the likelihood profile, using the confint function in lme4) for terms presented in bold do not cross 0. Samples with the same longitude and latitude (rounded to two decimal places) were grouped together under a random effects “site” term, to eliminate spatial autocorrelation (n = 253 samples, 95 groups). The resulting model residuals were not spatially autocorrelated (Moran’s I: observed = −.10, expected = −0.008, sd = 0.06, *p* = 0.12). A model excluding a term indicating sympatry with *C. virgo* provided a better fit to the data than the full model presented here (AIC_[model with ‘sympatric’]_ – AIC_[model without ‘sympatric’]_ = 3.98).

| model term | estimate | confidence int. |  |
| --- | --- | --- | --- |
|  |  | lower | upper |
| **intercept** | **-0.68** | **-0.73** | **-0.63** |
| site type (sympatric) | -0.07 | -0.15 | 0.02 |
| **Julian date** | **0.35** | **0.14** | **0.57** |
| **(Julian date)<sup>2</sup>** | **-0.31** | **-0.52** | **-0.10** |
| **latitude** | **-0.06** | **-0.09** | **-0.02** |
| longitude | 0.01 | -0.02 | 0.04 |

## Discussion

In a relatively brief period (one flight season for the demoiselle project and two for the smoky rubyspot project), we were able to compile a geographically and temporally diverse collection of photographs uploaded by participants to iNaturalist. With an increasing overall observation rate (Irwin 2018, Fig. 2a,b), it is likely that iNaturalist will continue to be an important resource for professional scientists and amateur naturalists alike. Though the user base of the iNaturalist website contributed the majority of observations, we found that our efforts to recruit participants through Twitter and the British Dragonfly Society did increase the relative number of new users contributing observations (Fig. 2c). Other websites, such as iSpot (Silvertown et al. 2015) and eBird (Sullivan et al. 2014), or Google images (Leighton et al. 2016), may also be useful platforms for generating photographic datasets.

Our pipeline for measuring citizen scientists’ photographs with a custom script in ImageJ (Supplementary material Appendix 1, 2) generated measurements of the proportion of wing surface pigmented that are highly correlated with measurements made under standardised conditions (Fig. 1). Although we used this script to measure the relative level of pigmentation in male damselfly wings, the script could easily be applied to a wide array of phenotypic measurements, such as measuring the relative size of different morphological features (e.g., eye size relative to head size (Werner and Seifan 2006), horn size relative to body size (Nijhout and Emlen 1998), relative wing spot size in butterflies (Monteiro et al. 1994)). Another recently developed package of tools for ImageJ could also be useful for extracting data on other aspects of patterning from citizen scientists’ photographs (Chan et al. 2019).

Previous research documented a seasonal increase in wing pigmentation, from the early season to the peak breeding season, in smoky rubyspots (Drury et al. 2015). Our analysis of the citizen scientist photos shows that male wing pigmentation decreases back to early season levels in the autumn. Moreover, analyses of geographical variation in the seasonal polyphenism of smoky rubyspot wing pigmentation demonstrate that, while seasonal shifts occur throughout the range, the magnitude of these shifts is greater in the Atlantic region (Figs. 3, A2), where individuals emerging in the peak breeding season can have completely melanised hindwings. Thus, using citizen scientist data has revealed that the seasonal and geographic patterns of variation in the wing coloration of this species are more complex than previously known.

In contrast to a study based on a traditional scientific sampling regime (Honkavaara et al. 2011), our analysis of citizen scientist photos collected at a higher spatial and temporal resolution (Figs. 4, A1) revealed no evidence for character displacement in male banded demoiselles’ wing pigmentation in Great Britain. After controlling for the effect of latitude, longitude, and observation date, sympatry was not a useful predictor of relative male wing spot area (Table 2). Another recent study in Fennoscandia also failed to find evidence that banded demoiselles in sympatry with beautiful demoiselles have smaller wing spots (in fact, they found the opposite pattern (Suhonen et al. 2018)). In contrast to these results about character displacement, we did find evidence to support previous research establishing an impact of latitude (Hassall and Thompson 2009) and observation date (Upton et al. 2016) on the relative area of male wing spots.

Although collaborating with the network of citizen scientists on iNaturalist.org is a powerful way to increase the spatial and temporal resolution of phenotypic sampling, such an approach is not free from bias (Boakes et al. 2010). In terms of spatial biases, some regions might be more highly visited than others, which could influence inferences made on patterns. Here, we found no impact of variation in observation rate on our inferences about phenotypes (Tables A6 & A9). Furthermore, we included a random effect for the grid cell in which observations were made in our models (Kuussaari and Helio 2007, Roy et al. 2012) to account for spatial autocorrelation in our models (e.g., Tables 2, 3). Temporal biases might also impact analyses, especially given the rapid increase in the number of observations available online. Here, we accounted for such temporal biases using a benchmark taxa approach (Hill 2012) in our analyses of observation accrual. There may also be biases in the individuals or species that are deemed worthy of noticing and photographing, which might be more problematic for some systems (e.g., rare or not commonly known species) and research questions than others. Comparisons of data from citizen scientists’ observations and systematically collected data like the ones we conducted here (Fig. 1) are one useful way to quantify the confidence that investigators can have in datasets collected by citizen scientists.

While iNaturalist is a useful tool for rapidly assembling a large number of observations from amateur naturalists, the most impactful citizen science collaborations are ones that engage with the community of citizen scientists throughout the life of the project (Pandya 2012). Moving the projects described here from being led in a top-down fashion toward a more collaborative model (sensu Tweddle, Robinson, Pocock, & Roy, 2012) with continued feedback and engagement would likely generate longer lasting citizen scientist engagement with and dedication to the projects (van der Wal et al. 2016), while simultaneously increasing community awareness of local organisms and their environments. Careful development of such mutually beneficial collaborations have the potential to open the door to statistically powerful analyses of phenotypic variation at unprecedented scales.

## Acknowledgements

We thank J. Fisher and E. Long for field assistance, C. Howard for statistical assistance, E. Colvey, D. Hepper, and F. McKenna at the British Dragonfly Society for help enlisting members, M. Cowen, S. McEachin, and P. Kong for feedback on the manuscript, and the many citizen scientists who participated in the project.

## Author Contributions

JPD & GFG designed the study, JPD, MB, & MH conducted fieldwork and curated the iNaturalist project pages, JPD, AF, & GFG conducted statistical analyses, JPD wrote the first draft of the manuscript, and all authors contributed to the final manuscript draft.

