## Supplementary material for "Continent-scale phenotype mapping using citizen scientists’ photographs"

Supporting Material for “**Continent-scale phenotype mapping using citizen scientists’ photographs”**

Appendix 1-2

Tables A1-A9

Figure A1-A2

**Appendix 1.** Instructions for making measurements on iNaturalist photos using ImageJ

• Download and install ImageJ from <https://imagej.nih.gov/ij/download.html>

• Opening images: Drag JPEG file onto the ImageJ icon on the sidebar.

• Optimizing view: You can re-size the image at any time to make better measurements; from the keyboard, use + to enlarge or - to shrink. To reposition the image, use the space bar to select the hand tool.

• Preparing macro: Copy and paste the macro script provided in Appendix S2 into a blank text file. Save macro file as ‘photograph_macro.txt’.

• Loading measurement tools: Go to Plugins > Macros > Install in the pulldown menu and open ‘photograph_macro.txt’ The macros will then be listed under Plugins > Macros, but you can run them by pressing the numbers shown.

First, measure the entire available wing surface:

• Setting the (arbitrary) scale: Before you can make area measurements, you need to set the scale of the image. For perching photos, there is no scale bar to use, so as a workaround, just use the line tool to draw a line across the individual’s eye. Then, press 1 (here and below, the number must be entered on the upper keyboard strip and not in the number keypad to the right) and then 1 again to set the scale of the image.

• Use the polygon tool to make area measurements 10 (the entire outline of all wings visible) and 11 (the entire outline of the wing spot [the basal wingspot in titia]) and 12 (the tip spot in titia). After you outline the area, press 1 and then enter the measurement code (see diagram).

• To change the code of a measurement, press 2 and then enter the number of the measurement and the new code.

• If you make a bad measurement, use the re-coding tool [2] to change the code of the bad measurement to 9 to tag it for deletion in excel. Then remeasure the area.

• Once you have taken all of the measurements, press 9. This copies the results to the clipboard for transfer to a datasheet and saves the analysed image file with a "z" before the name. IMPORTANT: To this file name, add a ‘z’ to the end of the image file, so that this photo is saved as ‘zz’.

Second, measure the hindwing surface only:

• Re-open the original image file, and repeat the procedure on the HINDWING only (1, then 10 [the outline of the entire hindwing surface], then 11 [the wingspot], and 12 [if present]), then save. When you finish and save this photo, there will be two new images corresponding to this file: 'z' (hw measured) and 'zz (whole wing surface measured)’ files for each photo.

**Appendix 2.** Macro script written for measuring photographs in ImageJ

```
//Global variables
var n = 1;
var scale = 0;
var length = 0;

macro "TAKE MEASUREMENT [1]" {
 	type = selectionType();
  	if (type==-1)
      exit("You need to select something first.");
      code = getNumber("code:", code);
	if (code>23)
      	exit("Invalid code. This version of the macro recognizes codes 1-23.");
	if (code<1)
      	exit("Invalid code. This version of the macro recognizes codes 1-23.");
	if (code==1) {//set scale
	//so that Measure yields area and xy:
	if (type!=5)
	      exit("Error: use line tool to set scale");
	run("Set Measurements...", "area centroid redirect=None decimal=5");
	run("Clear Results");
	n =0;
	getLine(x1, y1, x2, y2, lineWidth);
   	if (x1==-1)
      exit("Error: use line tool to set scale");
      scale = 2; // for a 2 mm scale line
	unit = "mm";
   	dx = x2-x1;
	dy = y2-y1;
   	length = sqrt(dx*dx+dy*dy);
	run("Set Scale...", "distance="+length+" known=scale pixel=0.99613 unit="+unit);
	// REM to set pixel to pixel aspect ratio for monitor
}
	if (scale==0)
      	exit("Need to set scale first, using 2 mm on scale bar.");
	if (code!=1 && type!=2)
		exit("Use polygon tool to measure areas.");
	run("Measure");
	setResult("Scale", n, length);
	setResult("Code", n, code);
	setResult("Image", n, getTitle);
	n = n+1;
	run("Overlay Options...", "stroke=yellow width=5 fill=none set");
	run("Labels...", "color=white font=14 show bold");
	run("Add Selection...");
}

macro "RECODE [2]" {
rn = getNumber("Measurement you want to recode (esc to cancel):", n);
if (rn==1)
      	exit("You can't recode the scale line.");
code = getNumber("New code:", code);
if (code>23)
      	exit("Invalid code. This version of the macro recognizes codes 1-23.");
if (code<1)
      	exit("Invalid code. This version of the macro recognizes codes 1-23.");
if (code==1)
	exit("You can't recode a measurement with the scale line code.");
setResult("Code", rn-1, code);  //rn-1 because first measurement is n = 0
}

macro "COPY RESULTS AND SAVE IMAGE [9]" {
String.copyResults
getDateAndTime(year, month, dayOfWeek, dayOfMonth, hour, minute, second, msec);
mo = month+1 // because Jan = 0
timestring = "Measurement date: " + year +" "+ mo +" "+ dayOfMonth;
setFont("SansSerif", 112, " antialiased");
setColor("black");
drawString(timestring, 102, 166);
//makeText(timestring, 102, 166);
path =  getDirectory("image");
zfile =  "z" + getTitle;
saveAs("Jpeg",  path + zfile);
```

}

**Table A1.** Study locations where standard and perching photos were taken for proof-of-principle analyses.

| **species** | **site code** | **latitude** | **longitude** |
| --- | --- | --- | --- |
| *H. titia* | LCPC1 | 31.2792 | -89.7131 |
|  | BF17 | 31.0749 | -93.4959 |
|  | NPKB1 | 31.5166 | -93.1433 |
|  | LCSB3 | 31.3222 | -89.7011 |
|  | LCBC4 | 31.2756 | -89.7311 |
|  | WCKC1 | 31.2692 | -90.1692 |
|  | PCCC1 | 30.988 | -89.0057 |
|  | TPAR1 | 30.7231 | -90.1408 |
| *C. splendens* | YODE01 | 53.84 | -0.946 |
|  | LIID01 | 53.457 | -0.8484 |
|  | NOID01 | 53.3629 | -0.9289 |
|  | NHWE01 | 52.519 | -0.7331 |
|  | NHIS01 | 52.4123 | -0.693 |
|  | SHRT01 | 52.3094 | -2.6662 |
|  | CATS01 | 52.3258 | -0.1097 |
|  | NHRT01 | 52.1324 | -0.9603 |
|  | CDRW01 | 54.7657 | -1.5542 |

**Table A2.** Linear regressions, modelling indices of relative wing pigmentation as measured from standard photographs (Y) as a function of indices measured on perched photographs (X).

| **species** | **measurement** | **model term** | **estimate** | **SE** | **t-value** | **p-value** |
| --- | --- | --- | --- | --- | --- | --- |
| *H. titia* | hindwing | intercept | -0.05 | 0.02 | -3.12 | 0.003 |
|  |  | X | 1.02 | 0.02 | 48.53 | < 0.001 |
|  |  |  |  |  | *Adjusted R^2^:* | 0.97 |
|  | entire wing | intercept | -0.06 | 0.02 | -3.92 | < 0.001 |
|  |  | X | 0.74 | 0.02 | 34.34 | < 0.001 |
|  |  |  |  |  | *Adjusted R^2^:* | 0.95 |
| *C. splendens* | hindwing | intercept | -0.14 | 0.02 | -6.12 | < 0.001 |
|  |  | X | 1.14 | 0.04 | 28.15 | < 0.001 |
|  |  |  |  |  | *Adjusted R^2^:* | 0.89 |
|  | entire wing | intercept | -0.13 | 0.02 | -5.75 | < 0.001 |
|  |  | X | 1.22 | 0.04 | 29.12 | < 0.001 |
|  |  |  |  |  | *Adjusted R^2^:* | 0.90 |

**Table A3.** Spearman rank correlations measuring inter-observer reliability in different measurement types.

| **species** | **photo** | **hindwing *ρ* (n)** | **entire wing *ρ* (n)** |
| --- | --- | --- | --- |
| *H. titia* | standard | 0.98 (32) | 0.99 (32) |
|  | perching | 0.98 (27) | 0.97 (28) |
| *C. splendens* | standard | 0.99 (29) | 0.98 (29) |
|  | perching | 0.88 (36) | 0.94 (35) |

**Table A4.** Mixed-effect regression of the proportion of the visible wing surface with pigment in *H. titia* males. The dependent variable was log-transformed, and longitude, latitude, julian date and julian date were standardized to a mean of zero and variance of 1, prior to analysis. Samples with the same longitude and latitude (rounded to two decimal places) were grouped together under a random effects “site” term, to eliminate spatial autocorrelation (n = 426 samples, 266 groups), the absence of which was verified with Moran’s I (observed = -0.0149, expected = -0.0027, sd = 0.0309, *p* = 0.69). The 95% confidence intervals (calculated from the likelihood profile, using the confint function in lme4) for terms presented in bold do not cross 0 (i.e., in this model, no terms’ confidence intervals crossed 0). A model excluding regional groupings provided a far worse fit to the data than the model presented here (AIC_[model with ‘region’]_ – AIC_[model without ‘region’]_ = -91.22). Tukey’s method was used for the multiple comparisons between regions.

**a. fixed effects**

| **model term** | **estimate** | **confidence int.** | |
| --- | --- | --- | --- |
|  |  | **lower** | **upper** |
| **intercept** | **-0.57** | **-0.63** | **-0.52** |
| **Northern region** | **-0.96** | **-1.17** | **-0.75** |
| **Pacific region** | **-0.48** | **-0.73** | **-0.24** |
| **Julian date** | **1.34** | **1.16** | **1.54** |
| **(Julian date)^2^** | **-1.49** | **-1.68** | **-1.30** |
| **latitude** | **-0.09** | **-0.15** | **-0.03** |
| **longitude** | **-0.08** | **-0.15** | **-0.01** |

**b. multiple comparisons**

| **contrast** | **estimate** | **std. error** | ***z*-value** | ***p*-value** |
| --- | --- | --- | --- | --- |
| Northern-Atlantic | -0.96 | 0.11 | -8.73 | <0.001 |
| Pacific–Atlantic | -0.48 | 0.13 | -3.81 | <0.001 |
| Pacific-Northern | 0.48 | 0.19 | 2.49 | 0.029 |

**Table A5.** Mixed-effect regression of the log-transformed proportion of the hindwing surface with pigment for *H. titia* males (*n* = 437), (a) excluding the ‘region’ variable from the main text and (b) including the Pacific/Atlantic regions, but lumping “northern” observations into the “Atlantic” region. Julian date and its quadratic term, latitude, and longitude were all z-transformed prior to model fitting. A ‘site’ variable was included as a random effect (see Table 2 caption). The 95% confidence intervals (calculated from the likelihood profile, using the confint function in lme4) for terms presented in bold do not cross 0 (i.e., in these models, no terms’ confidence intervals crossed 0). ΔAIC values presented are calculated as AIC_[model with ‘region’]_ – AIC_[model]_ , where the model with ‘region’ refers to the model presented in Table 2.

| **model** | **model term** | **estimate** | **confidence int.** | |
| --- | --- | --- | --- | --- |
|  |  |  | **lower** | **upper** |
| *(a) no region* | **intercept** | **-0.85** | **-0.91** | **-0.79** |
| ΔAIC = -91.63 | **Julian date** | **1.79** | **1.56** | **2.02** |
|  | **(Julian date)^2^** | **-1.94** | **-2.17** | **-1.72** |
|  | **latitude** | **-0.21** | **-0.27** | **-0.15** |
|  | **longitude** | **-0.36** | **-0.42** | **-0.29** |
| *(b) Pacific/Atlantic regions* | **intercept** | **-0.81** | **-0.87** | **-0.76** |
|  | **Pacific region** | **-0.86** | **-1.16** | **-0.55** |
| ΔAIC = -68.09 | **Julian date** | **1.56** | **1.32** | **1.80** |
|  | **(Julian date)^2^** | **-1.74** | **-1.97** | **-1.51** |
|  | **latitude** | **-0.28** | **-0.34** | **-0.21** |
|  | **longitude** | **-0.33** | **-0.39** | **-0.26** |

**Table A6.** Mixed-effect regression of the log-transformed proportion of the hindwing surface with pigment for *H. titia* males, binning observations for each month to test for an effect of observation rate on inferences (*n* = 315 site-by-month observations). Month, latitude, longitude, and the number of observations were all z-transformed prior to model fitting. A ‘site’ variable was included as a random effect (see Table 2 caption). The 95% confidence intervals (calculated from the likelihood profile, using the confint function in lme4) for terms presented in bold do not cross 0. A model excluding the region term provides a poorer fit, confirming an impact of region (AIC_[model with ‘region’]_ – AIC_[model without ‘region’]_  = -110.43).

| **model term** | **estimate** | **confidence int.** | |
| --- | --- | --- | --- |
|  |  | **lower** | **upper** |
| **intercept** | **-0.55** | **-0.63** | **-0.47** |
| **Northern region** | **-1.09** | **-1.40** | **-0.78** |
| **Pacific region** | **-1.32** | **-1.65** | **-0.99** |
| **month** | **-0.18** | **-0.25** | **-0.11** |
| latitude | -0.08 | -0.18 | 0.01 |
| longitude | -0.06 | -0.16 | 0.03 |
| observation count | 0.02 | -0.04 | 0.08 |

**Table A7.** Mixed-effect regression of the log-transformed proportion of the hindwing surface with pigment for *C. splendens* males (*n* = 253). Julian date and its quadratic term, latitude, and longitude were all z-transformed prior to model fitting. A ‘site’ variable was included as a random effect (*n* =253 observations, 95 groups, see Table 3 caption). Model residuals were not spatially autocorrelated (Moran’s I: observed = -0.12, expected = -0.008, sd=0.06, *p* = 0.06). The 95% confidence intervals (calculated from the likelihood profile, using the confint function in lme4) for terms presented in bold do not cross 0. A model excluding a term indicating sympatry with *C. virgo* provided a better fit to the data than the full model presented here (AIC_[model with ‘sympatric’]_ – AIC_[model without ‘sympatric’]_ = 4.37).

| **model term** | **estimate** | **confidence int.** | |
| --- | --- | --- | --- |
|  |  | **lower** | **upper** |
| **intercept** | **-0.59** | **-0.64** | **-0.55** |
| site type (sympatric) | -0.06 | -0.14 | 0.02 |
| **Julian date** | **0.52** | **0.21** | **0.64** |
| **(Julian date)^2^** | **-0.38** | **-0.59** | **-0.17** |
| **latitude** | **-0.05** | **-0.09** | **-0.02** |
| longitude | 0.02 | -0.01 | 0.05 |

**Table A8.** Mixed-effect regression of the log-transformed proportion of the visible wing surface with pigment for *C. splendens* males (*n* = 253), using two additional cut-offs for sympatry. Julian date and its quadratic term, latitude, and longitude were all z-transformed prior to model fitting. A ‘site’ variable was included as a random effect (see Table 3 caption). The 95% confidence intervals (calculated from the likelihood profile, using the confint function in lme4) for terms presented in bold do not cross 0. ΔAIC values presented compare presented model with a model excluding a term indicating sympatry with *C. virgo* (i.e., AIC_[model with ‘sympatric’]_ – AIC_[model without ‘sympatric’]_)

| **model** | **model term** | **estimate** | **confidence int.** | |
| --- | --- | --- | --- | --- |
|  |  |  | **lower** | **upper** |
| *conservative sympatry cut-off (one heterospecific reported within 2km, five reported within 20k)*  ΔAIC = 6.16 | **intercept** | **-0.71** | **-0.74** | **-0.68** |
|  | site type (sympatric) | 0.03 | -0.04 | 0.10 |
|  | **Julian date** | **0.36** | **0.15** | **0.58** |
|  | **(Julian date)^2^** | **-0.32** | **-0.53** | **-0.11** |
|  | **latitude** | **-0.04** | **-0.06** | **-0.01** |
|  | **longitude** | **0.03** | **0.01** | **0.06** |
| *very conservative sympatry cut-off (one heterospecific reported within 1km, five reported within 10k)*  ΔAIC = 5.69 | **intercept** | **-0.71** | **-0.74** | **-0.68** |
|  | site type (sympatric) | -0.02 | -0.14 | 0.09 |
|  | **Julian date** | **0.36** | **0.15** | **0.58** |
|  | **(Julian date)^2^** | **-0.32** | **-0.53** | **-0.11** |
|  | **latitude** | **-0.04** | **-0.07** | **-0.01** |
|  | **longitude** | **0.03** | **0.004** | **0.05** |

**Table A9.** Mixed-effect regression of the log-transformed proportion of the visible wing surface with pigment for *C. splendens* males, binning observations for each month to test for an effect of observation rate on inferences (*n* = 103 site-by-month observations). Month, latitude, longitude, and the number of observations were all z-transformed prior to model fitting. A ‘site’ variable was included as a random effect (see Table 3 caption). The 95% confidence intervals (calculated from the likelihood profile, using the confint function in lme4) for terms presented in bold do not cross 0. A model excluding the sympatry term provides a better fit, confirming a lack of an impact of sympatry with *C. virgo* (AIC_[model with ‘sympatric’]_ – AIC_[model without ‘sympatric’]_  = 3.27).

| **model term** | **estimate** | **confidence int.** | |
| --- | --- | --- | --- |
|  |  | **lower** | **upper** |
| **intercept** | **-0.65** | **-0.71** | **-0.60** |
| site type (sympatric) | -0.08 | -0.17 | 0.01 |
| **month** | **0.05** | **0.02** | **0.07** |
| **latitude** | **-0.07** | **-0.11** | **-0.03** |
| longitude | 0.02 | -0.02 | 0.06 |
| observation count | 0.03 | -0.002 | 0.06 |

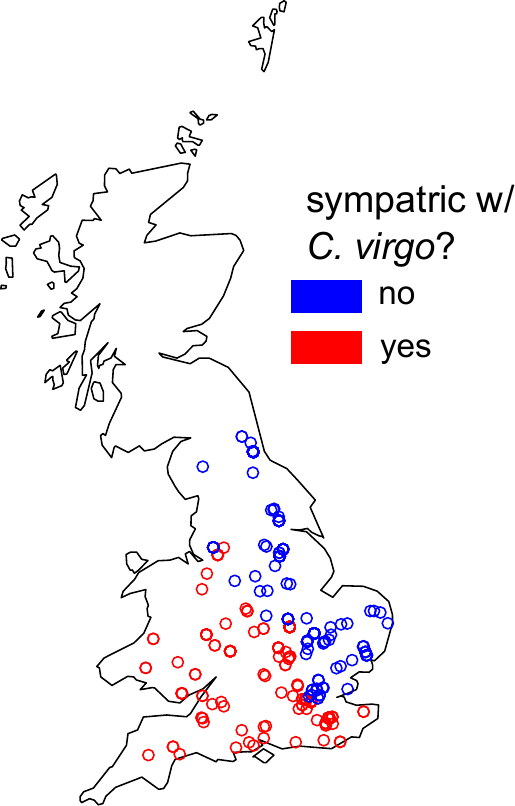

Figure A1. Map of *Calopteryx splendens* observations in Great Britain and their designation as sympatric or allopatric.

Figure A2. Predicted levels of wing pigmentation in the early season (Julian date = 75), peak season (Julian date = 200) and late season (Julian date = 325) from each of the three models with different region categories. Predictions across all models are largely concordant, but a model with discrete regions for Atlantic, Pacific, and northern populations (top) greatly outperforms models with only latitude and longitude (middle) or a model that lumps Atlantic and northern populations (bottom). Predicted values larger than 1 were set to 1 for plotting.
